# Genome-wide identification of accessible chromatin regions in bumblebee (*Bombus terrestris*) by ATAC-seq

**DOI:** 10.1101/818211

**Authors:** Xiaomeng Zhao, Weilin Xu, Sarah Schaack, Cheng Sun

**Affiliations:** Institute of Apicultural Research, Chinese Academy of Agricultural Sciences, Beijing, China; Department of Biology, Reed College, Portland, OR, USA

**Keywords:** bumblebee, *Bombus terrestris*, ATAC-seq, accessible chromatin region, regulatory element, developmental stage, metamorphosis

## Abstract

Bumblebees (Hymenoptera: Apidae) are important pollinating insects that play pivotal roles in crop production and natural ecosystem services. To date, while the protein-coding sequences of bumblebees have been extensively annotated, regulatory elements, such as promoters and enhancers, have been poorly annotated in the bumblebee genome. To achieve a comprehensive profile of accessible chromatin regions and provide clues for all possible regulatory elements in the bumblebee genome, we did ATAC-seq (Assay for Transposase-Accessible Chromatin with high-throughput sequencing) for *B. terrestris* samples derived from its four developmental stages: egg, larva, pupa, and adult, respectively. The sequencing reads of ATAC-seq were mapped to *B. terrestris* reference genome, and the accessible chromatin regions of bumblebee were identified and characterized by using bioinformatic methods. Our study will provide important resources not only for uncovering regulatory elements in the bumblebee genome, but also for expanding our understanding of bumblebee biology. The ATAC-seq data generated in this study has been deposited in NCBI GEO (accession#: GSE131063).

## Introduction

Regulatory elements play a major role in controlling the temporal and spatial expression of genes, through which they control the development and physiology of an organism (Narlikar and Ovcharenko 2009). Mutations that affect the function of regulatory sequences contribute to phenotypic diversity within and between species, and can also lead to disease phenotypes or change an organism’s susceptibility to disease (Hardison and Taylor 2012; Wittkopp and Kalay 2012). Due to the increasing awareness of the importance of regulatory elements (Gallagher and Chen-Plotkin 2018), more and more tools are available to identify such sequences genome-wide (Hardison and Taylor 2012).

Bumblebees (Hymenoptera: Apidae) are important pollinating insects that play pivotal roles in crop production and natural ecosystem services (Fontaine et al. 2006; Garibaldi et al. 2013; Velthuis and van Doorn 2006). Also, they are holometabolous insects that undergo four developmental stages (egg, larva, pupa, adult), so they are useful models to study mechanisms underlying developmental plasticity (Tian and Hines 2018). To date, the protein-coding sequences of bumblebees have been extensively annotated (Sadd et al. 2015). However, regulatory elements, such as promoters, enhancers and silencers etc., have been poorly annotated in the bumblebee genome.

ATAC-seq (Assay for Transposase-Accessible Chromatin with high-throughput sequencing) is a newly developed method that can determine accessible chromatin regions across the genome (Buenrostro et al. 2013; Buenrostro et al. 2015), from which regulatory elements could be inferred. This technique not only requires less starting material, but also produces more precise results than previous approaches (Tsompana and Buck 2014; Lai and Pugh 2017). In addition, ATAC-seq has demonstrated its power to detect chromatin accessibility using whole animal preparations (containing mixtures of tissues or organs) with high sensitivity (Daugherty et al. 2017).

In this study, we used ATAC-seq to do a genome-wide survey of accessible chromatin regions in *Bombus terrestris*, one of the most widely used commercial bumblebee species world-wide (Velthuis and van Doorn 2006). To achieve a comprehensive profile of open chromatin regions and provide clues for all possible regulatory elements in bumblebee genome, we did ATAC-seq for *B. terrestris* samples derived from its four developmental stages: egg, larva, pupa, and adult, respectively. The accessible chromatin regions identified by this study will provide important resources for uncovering promoters, enhancers and other regulatory elements in the bumblebee genome. In addition, the obtained information will expand our understanding of bumblebee biology and facilitate the cloning of bumblebee genes that control important traits.

## Materials and Methods

### Bumblebee samples for ATAC-seq

Commercial *B. terrestris* colonies were bought from Koppert China (http://www.koppert.cn). Worker bee samples were collected from each of the four developmental stages: egg, larva, pupa, and adult, respectively, and samples from each developmental stage were put into one 1.5 ml of centrifuge tube to ensure the tissue sampled was equivalent to about the size of one adult worker bee. The eggs we collected were straight and smooth; all larvae had a C-shape curve; pupae had visible compound eye pigmentation and clear head-thorax-abdomen segmentation, but their wings were not developed yet; adult bees were bright with dense hair, and could flap their wings. All samples were flash frozen after tissue collection.

### ATAC-seq protocol

ATAC-seq was performed by BGI-Shenzhen (https://en.genomics.cn). Briefly, count and harvest about 50,000 intact and homogenous cells for each developmental stage, which were then centrifuged for 5min at 500 ×g, 4°C; After discarding supernatant, the pellet was gently re-suspended with 50 µL of cold 1x PBS buffer, which followed by centrifuging them again for 5min at 500 ×g, 4°C. After removing supernatant, the pellet was gently pipetted and resuspended in 50 µL of cold lysis buffer (10 mM Tris-HCl, pH 7.4, 10 mM NaCl, 3 mM MgCl_2_, 0.1% IGEPAL CA-630) to release nuclei. After lysis, the suspension was spun at 500 × g for 10 minutes, 4°C. After centrifugation, the pellet was immediately resuspended in the transposase reaction mix (25 μL 2x TD buffer, 2.5 μL Transposase (Illumina) and 22.5 μL of nuclease free water). The transposition reaction was carried out at 37 °C for 30 minutes. After incubation, the products were purified by Qiagen MinElute PCR Purification Kit, and transposed DNA was eluted in 10 µL Elution Buffer (10mM Tris buffer, pH 8). The purified products were amplified in a 50 µL reactions containing the purified transposed DNA, 1x NEBnext High-Fidelity PCR master mix, and 1.25 μM of custom Nextera PCR primers, with the following PCR program: 1) 72°C, 5 minutes; 2) 98°C, 30 seconds; 3) 98°C, 10 seconds; 4) 63°C, 30 seconds; 5) 72°C, 1 minute; 6) Repeat steps 3-5 4 times; 7) Hold at 4°C. After amplification, the PCR products obtained were purified by Qiagen MinElute PCR Purification Kit, with the purified PCR products being eluted in 20 µL Elution Buffer (10mM Tris Buffer, pH 8). Next, the purified PCR products were used to produce single-strand DNA circles, from which DNA nanoballs were generated by rolling circle replication as previously described (Huang et al. 2017). Finally, the DNA nanoballs were sequenced on the BGISEQ-500 sequencing platform, generating paired-end reads with a read length of 50 bp.

### Sequence processing, mapping, peak calling and nucleosome positioning

Raw reads were filtered first to remove low-quality reads and adaptor sequences by SOAPnuke (Chen et al., 2018), with ‘filter -l 5 -q 0.5 -n 0.1 -Q 2 −5 1 -c 50’ parameters. Cleaned reads were mapped to the reference genome of *B. terrestris* (GenBank: GCF_000214255.1) using Bowtie2 (version: 2.2.5) (Langmead and Salzberg, 2012), with ‘-q --phred64 --sensitive --dpad 0 --gbar 99999999 --mp 1,1 --np 1 --score-min L,0,-0.1 -I 1 -X 1000 -p 16 -k 200’ parameters. The fragment length distribution of ATAC-seq was determined by the “fragSizeDist” function in the R package ATACseqQC (Ou et al. 2018). Sequencing depth distribution was obtained by BEDTools (version: 2.23.0) (Quinlan and Hall, 2010) with ‘genomecov’ command. We used MACS2 (version: 2.1.2) to call peaks (open chromatin regions) with ‘--nomodel --extsize 200 --shift −100 --format BAM --gsize 2.17e8 --call-summits’ parameters as used before (Zhang et al., 2008; Daugherty et al. 2017; Cusanovich et al. 2018). Peaks from different developmental stages were integrated and the profile of open chromatin signals around genes for each developmental stage was plotted by ChIPseeker (Version 1.18.0) (Yu et al., 2015). The relative position of peaks to the nearest protein-coding gene was annotated by comparing their coordinates with the annotation of *B. terrestris* genome, with the following priority order: promoter (−2kb, TSS), 5’UTR, 3’UTR, exon, intron, downstream (TES, 3kb) and distal intergenic region. To identify the position of nucleosomes, broad peaks called by MACS2 were analyzed using NucleoATAC (version: 0.3.4) (Schep et al., 2015).

## Results and Discussion

### ATAC-seq and sequence mapping

Fresh tissues were collected from the four developmental stages of *B. terrestris*, which were used to assay for transposase-accessible chromatin with high-throughput sequencing (ATAC-seq). A total of 153, 187, 216 and 128 million reads were generated for the four developmental stages of *B. terrestris*: egg, larva, pupa, and adult stage, respectively. After removal of adaptor sequences and low-quality reads, 152, 186, 216, and 127 million clean reads were obtained for egg, larva, pupa, and adult stage, respectively, with the total length of clean reads > 6 Gb for every developmental stage (Table 1). Clean reads obtained from the four developmental stages were mapped to the reference genome of *B. terrestris* using Bowtie2, with a total of 93, 80, 139, and 64 million reads could be uniquely mapped, respectively (Table 1). Based on the mapping results, we inferred the fragment size distribution of ATAC-seq data. From the results we could see that, as expected, while a majority of fragments were shorter than one nucleosome length (approximately 150 bp), there were significant number of fragments longer than this length (Figure 1A).

**Table 1.** The summary of ATAC-seq, read mapping and peak calling.

|  | Egg | Larva | Pupa | Adult |
| --- | --- | --- | --- | --- |
| Raw reads # (million) | 153 | 187 | 216 | 128 |
| Clean reads # (million) | 152 | 186 | 216 | 127 |
| Total length of clean reads (Gb) | 7.6 | 9.3 | 10.8 | 6.4 |
| Mapped reads # (million) | 126 | 119 | 173 | 78 |
| Uniquely mapped reads # (million) | 93 | 80 | 139 | 64 |
| Candidate peak # | 58,509 | 36,291 | 47,141 | 36,939 |
| Filtered peak # | 44,602 | 22,797 | 35,590 | 25,284 |
| Total length of candidate peak (Mb) | 25 | 16 | 22 | 16 |
| Total length of filtered peak (Mb) | 19 | 10 | 16 | 11 |
| Average length of candidate peak (bp) | 430 | 431 | 460 | 435 |
| Average length of filtered peak (bp) | 424 | 427 | 458 | 439 |
| Chromosome accessibility rate (%) | 8.01 | 4.12 | 6.90 | 4.69 |

Note: Egg, Larva, Pupa and Adult represent the four development stages of *B. terrestris*, respectively. Candidate peaks were called by MACS2 with p-value < 1e-1. Filtered peaks were future refined from candidate peaks to only keep peaks that were located on chromosomes and had a p-value < 1e-5.

**Figure 1.**
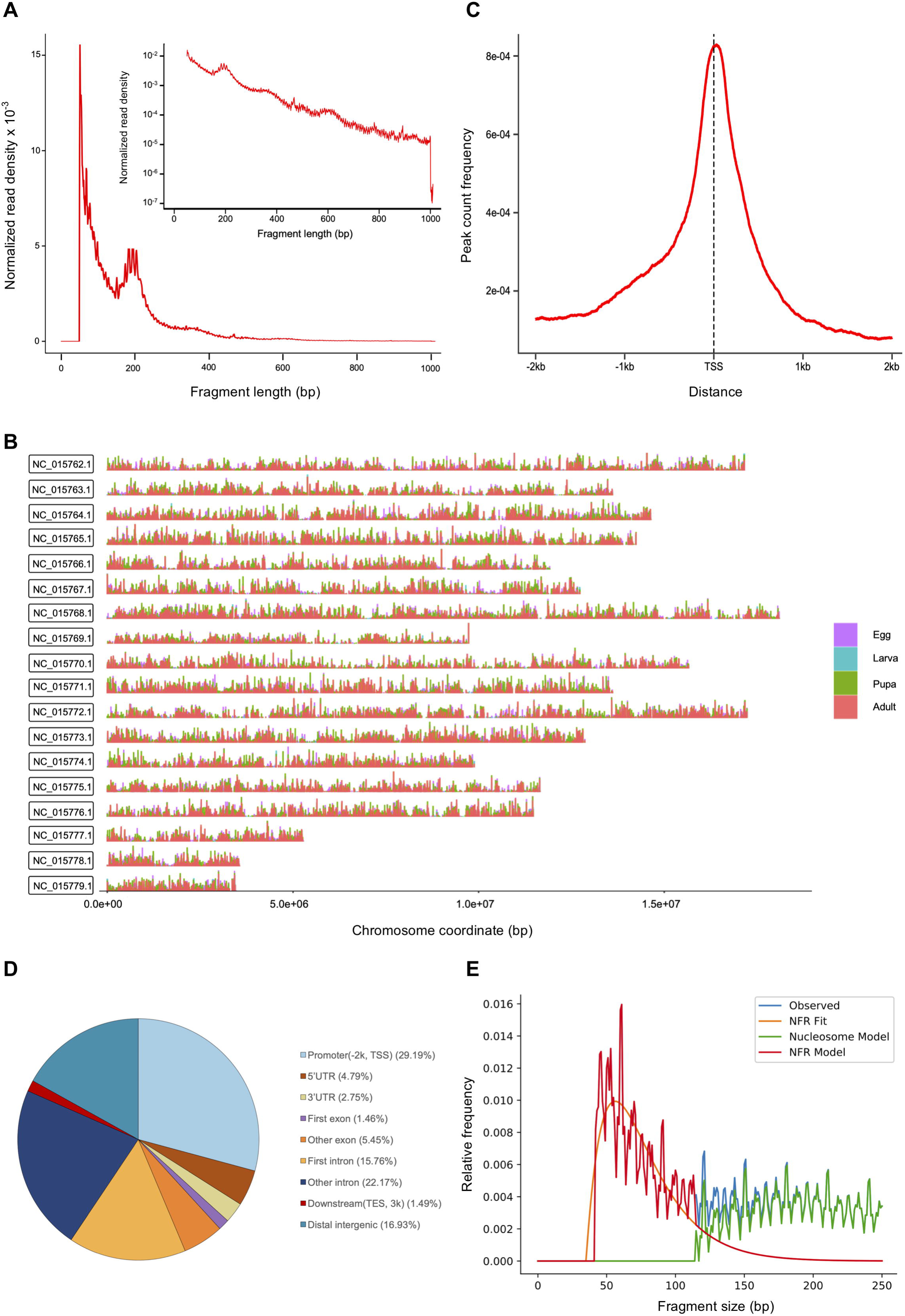
The features of ATAC-seq data for bumblebee. **(A)** Fragment size distribution of ATAC-seq data for the larval developmental stage. **(B)** Peak profiles obtained by integrating peaks from the four developmental stages showing a comprehensive picture of bumblebee chromatin accessibility. (**C**) The profile of chromatin accessible signals around bumblebee genes (based on the larval developmental stage). TSS represents transcription start sites. **(D)** The relative location of peaks relative to their nearest genes (based on the larval developmental stage). **(E)** Model for nucleosome positioning and NFR (nucleosome free region) distribution at the larval developmental stage predicted by NucleoATAC.

### Delimitation and characterization of accessible chromatin regions

For each developmental stage, peaks (accessible chromatin regions) were called by MACS2 software, and obtained peaks were future filtered to only keep peaks that were located on the assembled chromosomes of *B. terrestris* and had a p-value <1e-5. After filtering, 44,602, 22,797, 35,590 and 25,284 accessible chromatin regions were remained for developmental stage of egg, larva, pupa, and adult, respectively (Table 1). The average length of accessible chromatin regions is similar among the four developmental stages (ranges from 424 to 458 bp; Table 1). In total, 8.01, 4.12, 6.90 and 4.69 percent of *B. terrestris* genome comprises of accessible chromatin in each stage (egg, larva, pupa, and adult, respectively; Table 1).

The peak files from all four developmental stages have been deposited to the NCBI Gene Expression Omnibus (accession number: GSE131063), and can be uploaded to UCSC genome browser (http://genome.ucsc.edu/) individually to reveal the distribution of open chromatins across the genome in each specific developmental stage. The genome-wide distribution of all the potential accessible chromatin regions in the bumblebee genome was obtained by integrating identified accessible chromatin regions from all four developmental stages by using ChIPseeker (Figure 1B). In the figure, if more than one color present at a specific peak (accessible chromatin region), it indicates that the region could be detected as open chromatin in more than one developmental stage (Figure 1B).

The level of ATAC-seq signal corresponds to the level of chromatin accessibility (Buenrostro et al. 2013) and can be used to identify poised and active regulatory regions genome-wide (Daugherty et al. 2017). We plotted chromatin accessible signals around genes for each developmental stage, and the result for the larva stage is shown in Figure 1C. Notably, chromatin accessible signals are enriched at the transcription start sites (TSSs) of bumblebee genes.

To understand the relative position of accessible chromatin regions to their nearest gene, we compared the coordinates of peaks with that of *B. terrestris* protein-coding genes. From the results we could see that, if we consider 2 kb upstream of TSS (transcription start site) as promoter region, ~30% of accessible chromatin is located in promoter regions (Figure 1D). Open chromatin regions could also be found in exons, introns and distal intergenic regions (Figure 1D). Enhancers may be located far from their target genes or even within exons and introns (Birnbaum et al. 2012; Pennacchio et al. 2013), therefore, the identified open chromatin in such regions represent major candidates for future studies for enhancers in bumblebee genome.

### Nucleosome positioning analysis

Nucleosome positioning is also a major determinant of gene expression (Radman-Livaja and Rando 2009; Chereji and Clark 2018). We used NucleoATAC to predict the nucleosome position and the nucleosome free region (NFR) of *B. terrestris* in each of the four developmental stages and results have been deposited to the NCBI Gene Expression Omnibus (accession #: GSE131063). The result of nucleosome positioning for the larval developmental stage is summarized in Figure 1E.

While most genomic DNA is occupied by nucleosomes, many regulatory elements are depleted of nucleosomes (i.e., have low occupancy) (Struhl and Segal 2013). In addition, transcription factors may bind DNAs in NFRs, which would make Tn5 transposase hard to insert so that relatively lower ATAC signals might be obtained in those regions (Jiang and Pugh 2009; Buenrostro et al. 2013). Therefore, the nucleosome positioning information identified in this study will provide useful clues to identify regulatory elements and infer transcription factor binding sites in bumblebee genome.

## Conclusions

We have successfully generated ATAC-seq data for bumblebee samples derived from its four developmental stages: egg, larva, pupa, and adult, respectively. Using this dataset, all the possible accessible chromatin regions of bumblebee were identified and characterized. Our study will provide important resources not only for uncovering regulatory elements in the bumblebee genome, but also for expanding our understanding of bumblebee biology.

## Data access

The raw sequence data, as well as stage-specific peak files and nucleosome positioning files, have been deposited to the NCBI Gene Expression Omnibus (NCBI GEO) (Edgar et al. 2002) and are accessible through GEO Series accession number GSE131063.

## Author contributions

CS and XZ conceived the study. XZ performed bioinformatics analysis. WX and XZ collected samples. WX and XZ are both involved in ATAC-seq. CS, XZ and SS wrote the manuscript. All authors read and approved the final manuscript.

## Funding

This work was supported by the Elite Youth Program of Chinese Academy of Agricultural Sciences [to CS], and National Natural Science Foundation of China [31971397].

## Conflict of interests

The authors declare that the research was conducted in the absence of any commercial or financial relationships that could be construed as a potential conflict of interest.

